# Neuromodulation and Differential Learning Across Mosquito Species

**DOI:** 10.1101/755017

**Authors:** Gabriella H. Wolff, Chloé Lahondère, Clément Vinauger, Jeffrey A. Riffell

## Abstract

Mosquitoes can learn to change their host-feeding behaviors, such as shifting activity times to avoid bednets or switching from biting animals to biting humans, leading to the transfer of zoonotic diseases. Dopamine is critical for insect learning, but its role in the antennal lobe remains unclear, and it is unknown whether different mosquito species learn the same odor cues. We assayed aversive olfactory learning and dopaminergic brain innervation in four mosquito species with different host preferences and report here that they differentially learn odors salient to their preferred host and innervation patterns vary across species. Using genetically-encoded GCaMP6s *Aedes aegypti*, we mapped odor-evoked antennal lobe activity and report that glomeruli tuned to “learnable” odors have significantly higher dopaminergic innervation. Changes in dopamine expression in the antennal lobes of diverse invertebrate species may be an evolutionary mechanism to adapt olfactory learning circuitry without changing brain structure and for mosquitoes an ability to adapt to other hosts when their preferred are no longer present.

## Introduction

There are over 3500 species of mosquitoes that exhibit remarkable diversity in feeding preferences, from flowers to humans and other mammals, and even frogs or birds^1,2^. Despite their often specific preferences for blood-hosts, mosquitoes exhibit a high degree of behavioral flexibility^3^. One mechanism for this flexibility is their capacity to learn and modify feeding preferences. For example, *Aedes aegypti* mosquitoes can learn the association between an odor and an aversive stimulus, such as the near-encounter of a swat, although only certain odors can be learned in this association^4^. The neuromodulator dopamine is critical for this learning, with dopaminergic neurons innervating specific subunits called glomeruli in the mosquito antennal lobe (Extended Data Fig. 1a), possibly allowing enhanced neuromodulation and drive of glomeruli that encode specific learnable odorants. Nonetheless, it remains an open question whether other mosquito species show a similar aptitude for learning, or what is the relationship between the saliency of the learned odors and differential dopaminergic innervation.

## Results

### Differential learning behaviors across species

To assess learning behaviors in different mosquito species, we first examined the antennal responses of four mosquito species that differ in their host preferences: *Ae. aegypti, Anopheles stephensi, Toxorhynchites amboinensis* and *Culex quinquefasciatus*; as adults, females of these species prefer to feed on humans, birds, and flowers, respectively^2,5^. Antennal responses to three odorants 1-octen-3-ol, and hexanoic acid (both associated with vertebrate hosts), and linalool, a terpene alcohol produced by plants and a component of many floral scents^6–8^ - were recorded via electroantennogram (EAG), which is thought to measure the bulk sum of olfactory sensory neuron response on the antennae^9^. Although each species exhibited a significantly different pattern of responses to the odorants (Fig. 1; ANOVA: p<0.05), the three odorants consistently elicited robust responses across the different species. Together, these odorants allowed us to examine whether differential learning of odors is related to their host/feeding preferences.

**Fig. 1.**
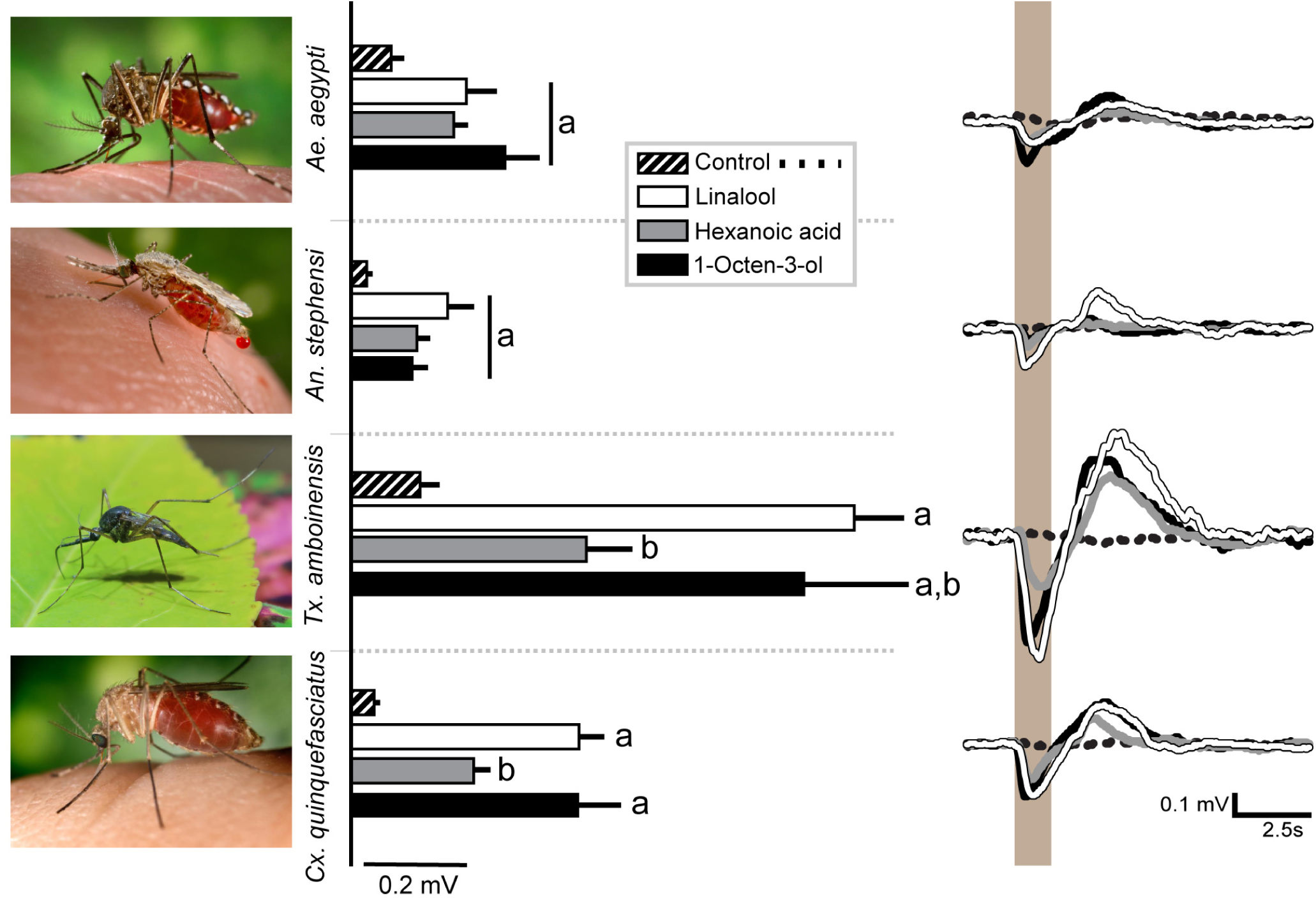
Antennal responses to host and plant odors vary across species. (Left) Four species were tested: *Ae. aegypti, An. stephensi, Tx. amboinensis*, and *Cx. quinquefasciatus*. (Middle) Histograms are the mean responses to odors grouped by species (mean ± SEM). For the strongly anthropophilic species, *Ae. aegypti* and *An. stephensi*, responses to all three odors were not statistically significant (p>0.05). By contrast, hexanoic acid elicited significantly lower responses compared to the other odorants in *Cx. quinquefasciatus* and linalool for *Tx. amboinensis* (t-test: p<0.05). Letters denote significant differences between odor stimuli (t-test: p<0.05). (Right) Average traces from 5 individuals of each species to the three odorants (1-octen-3-ol; linalool; hexanoic acid, and solvent (mineral oil) control.

Female mosquitoes from each species were trained in an aversive conditioning paradigm (Extended Data Fig. 1b, ^4^) using mechanical shock as the unconditioned stimulus (US) and one of the three odorants identified by EAG as the conditioned stimulus (CS). Although appetitive conditioning experiments conducted by our group also showed odorant-specific learning, in this study we used an aversive conditioning paradigm which allows use of the same unconditioned stimulus (US; a mechanical shock that mimics “swatting”) across species. Mosquitoes were divided into three treatment groups: “naïve” mosquitoes had no prior exposure to any test odors, “trained” mosquitoes were exposed to 10 pairings of the US and CS, and “unpaired” mosquitoes were handled in the same manner as trained mosquitoes, but odor and mechanical shock were presented randomly without temporal contingency. After 24 hours, mosquitoes were individually tested in a two-choice y-maze olfactometer and a preference index was calculated (Extended Data Fig. 1b). If the association was learned, we expected the preference index of trained mosquitoes to be significantly lower than the naïve and unpaired groups.

Training and testing of the different species to the odorants and mechanical shock showed that trained *Ae. aegypti* significantly avoided both hexanoic acid and 1-octen-3-ol compared to naïve (Fig. 2a, Binomial exact test, p=0.007 for 1-octen-3-ol and 0.0003 for hexanoic acid) or unpaired groups (Extended Data Fig. 2a). However, naïve and trained *Ae. aegypti* did not significantly differ in their preference for linalool (p=0.26). These results suggest that *Ae. aegypti* learned to avoid the odors associated with their preferred mammalian hosts, but not a common flower odorant. Like *Ae. aegypti, An. stephensi* also learned to avoid 1-octen-3-ol (Fig. 2b, p=0.0007). However, when hexanoic acid or linalool was used as the conditioned stimulus, trained *An. stephensi* exhibited lower preference indices compared to naïve mosquitoes, but values were not significantly different (p=0.22 for hexanoic acid and p=0.58 for linalool). By contrast, *Cx. quinquefasciatus*, which rarely feed on mammalian hosts and prefer avian hosts when they are available, did not show an ability to learn in this context (Fig. 2c). Previous work has shown that *Cx. quinquefasciatus*, unlike *Ae. aegypti* and *An. stephensi*, are repelled by 1-octen-3-ol which is found in mammalian, but not avian skin volatiles^10,11^. However, we did not observe significant differences in preference between naïve and trained groups for either 1-octen-3-ol, hexanoic acid, or linalool (Fig. 2c, p=0.93 for 1-octen-3-ol, p=0.83 for hexanoic acid, p=0.57 for linalool). We further trained a group of *Cx. quinquefasciatus* to nonanal, a major component of bird headspace profiles, which is an attractant for this species (Extended Data Fig. 2b; ^12^). Again, we did not observe evidence of learning in this species using our classical conditioning paradigm although activity levels were high, with 93-100% of *Cx. quinquefasciatus* making a choice in the olfactometer (Extended Data Table 1, p=0.91).

**Fig. 2.**
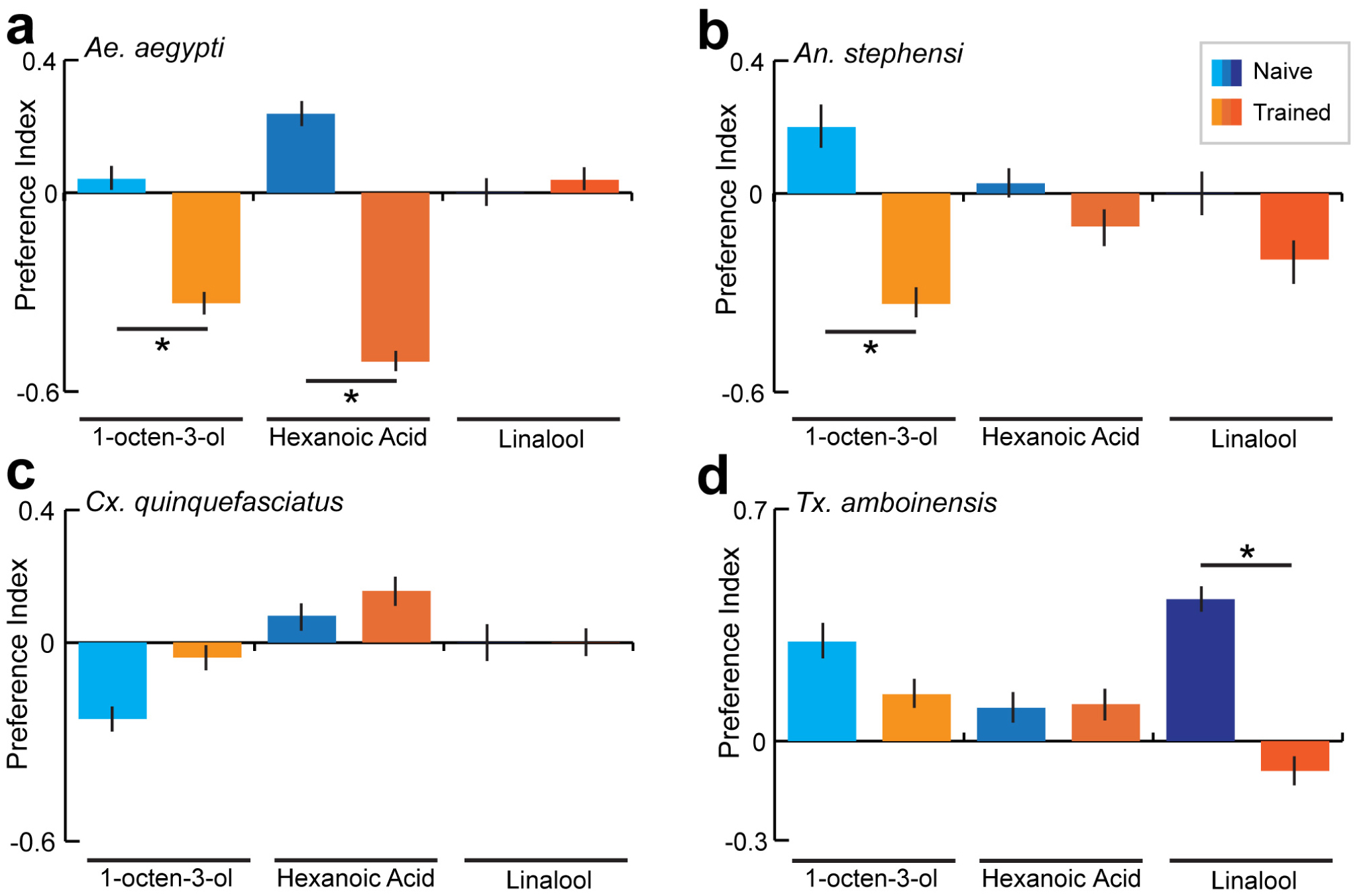
Mosquitoes learn odors associated with their preferred host. Mosquito preference index (PI) of (**a**) *Ae. aegypti*, (**b**) *An. stephensi*, (**c**) *Cx. quinquefasciatus*, and (**d**) *Tx. amboinensis* females to 1-octen-3-ol, hexanoic acid, and linalool in naïve and trained groups. Asterisks represent statistical significance from naïve PI (Binomial exact test: p<0.05). Bars are the mean ± SEM.

We next tested a nectar-feeding (non-hematophagous) mosquito, *Tx. amboinensis* to assess whether this species could learn associations with either of the vertebrate host or plant odors. When tested against 1-octen-3-ol or hexanoic acid, the trained mosquitoes did not differ significantly in preferences from the naïve groups (Fig. 2d) (Binomial exact test, p=0.20 for 1-octen-3-ol and p=0.75 for hexanoic acid). However, *Tx. amboinensis* was the only species in this study to learn the association between linalool and the mechanical shock (Binomial exact test, p=0.009), suggesting that like *Ae. aegypti* and *An. stephensi, Tx. amboinensis* learned the odor associated with their preferred food resource.

### Differential localization of TH-like immunoreactivity across species

The relationship between learning and dopaminergic innervation in the antennal lobe of mosquitoes remains unknown. In *Ae. aegypti*, there is strong, differential innervation in specific glomeruli and, in parallel, these mosquitoes can learn only specific odorants^4,13^. As a first step to examine whether differential learning across mosquitoes is associated with variations in dopamine innervation, we used antisera against tyrosine hydroxylase (TH), an enzyme required for the biosynthesis of dopamine, to localize this neuromodulator in the brain of each species. In particular, we focused on olfactory circuitry including the antennal lobes (ALs) which are the primary olfactory brain structures in insects, and the mushroom bodies (MBs) which are higher-order centers implicated in learning and memory (Extended Data Fig. 1a).

Although TH-like immunoreactivity was localized in the ALs and MBs of all four species, the degree of dopaminergic innervation varied dramatically. The most striking difference in TH-like immunoreactivity across mosquito species was differential innervation of the ALs and their substructures (Fig. 3a-d). Each AL is subdivided into spherical-shaped compartments of dense neuropil called olfactory glomeruli. Olfactory receptor neurons (ORNs) expressing the same ligand-gated receptor terminate convergently, generally in one glomerulus where they synapse onto projection neurons (PNs) that relay information to higher-order centers such as the MBs and lateral horns^14,15^. This organization compartmentalizes encoding of single odor chemicals into one or very few glomeruli, and the pattern of activity across glomeruli forms a combinatorial code designating an identity to complex odors ^16^. In *Ae. aegypti*, the ALs are extensively innervated with dopaminergic neurons, but TH-like immunoreactivity was heterogeneous across glomeruli (Fig. 3a,a’). The higher concentrations of TH were observed in the more anteromedial glomeruli, such as the AM1, AD2, MD1, MD2, and MD3 glomeruli^17^. Similar extensive but heterogenous dopaminergic innervation of the ALs was observed in *Tx. amboinensis*. However, the pattern of high- and low-concentration glomeruli was different in this species, with higher concentrations localized in posterolateral glomeruli (Fig. 3b,b’).

**Fig. 3.**
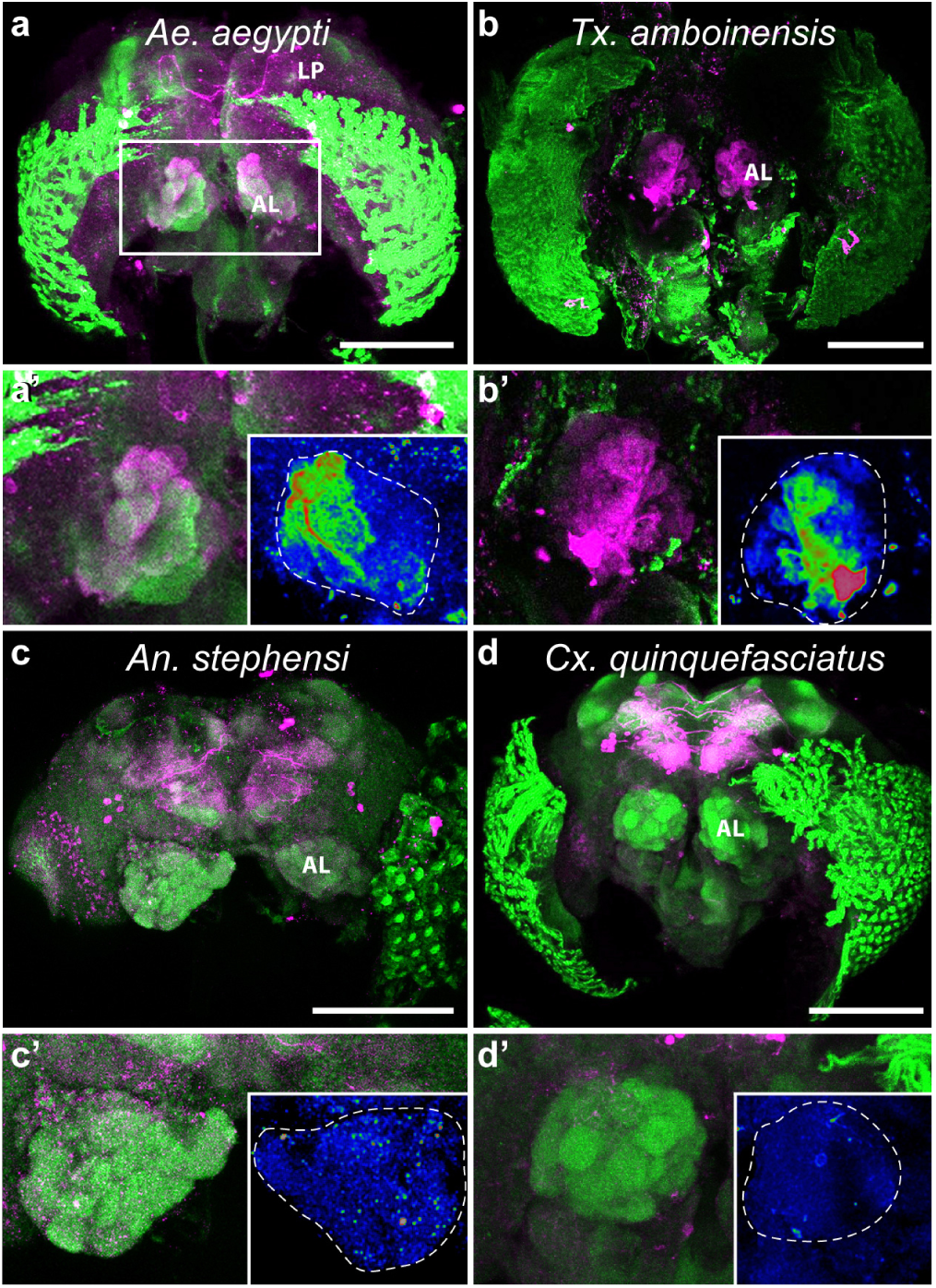
Differential dopaminergic innervation of antennal lobes across mosquito species. Confocal laser scans of **a**, *Ae. aegypti* ; **b**, *Tx. amboinensis*; **c**, *An. stephensi*; and **d**, *Cx. quinquefasciatus* female whole brains labeled with antisera against tyrosine hydroxylase to stain dopaminergic neurons (TH, magenta) and phosphorylated CaMKII (pCaMKII, green) for background staining. The box in A represents the areas magnified 2X in **a’-d’**. Insets: Tyrosine hydroxylase intensity pseudocolored using the Rainbow RGB lookup table in ImageJ. Dashed outlines delimitate AL boundaries. Scale bars = 100µm in **a**,**c**,**d** and 200µm in **b**. AL = antennal lobe, LP = lateral protocerebrum.

TH-like immunoreactivity signals were lower in the ALs of *An. stephensi*, but dopaminergic innervation was observed surrounding glomeruli, especially in the lateral region of the antennal lobes (Fig. 3c,c’). By contrast, in *Cx. quinquefasciatus* the TH signal was not detectable above background in the ALs although extensive TH-like immunoreactivity was observed in the lateral protocerebrum, surrounding the MBs in this species (Fig. 3d,d’). We used two different antisera, one raised against TH from rat (AB 572268) and one from Manduca sexta^18^ and both revealed the same pattern of immunoreactivity.

### Dopamine is necessary for olfactory aversive learning in *Ae. aegypti* and *An. stephensi*

The differences in dopaminergic innervation of ALs across mosquito species raises the question of the importance of dopamine for aversive learning in other mosquito species. To answer this question, we aversively trained two more groups of *An. stephensi* using 1-octen-3-ol as the conditioned stimulus: one group was injected with a dopamine antagonist (SCH-23398) and the other group was injected with physiological saline. As observed in *Ae. aegypti*, injection with the dopamine antagonist abolished aversive learning of 1-octen-3-ol in *An. stephensi* while saline-injected mosquitoes learned to avoid this odor (Fig. 4, ^4^). Despite differential dopaminergic innervation of ALs, both species required dopamine to form aversive associative memory of the host odor.

**Fig. 4.**
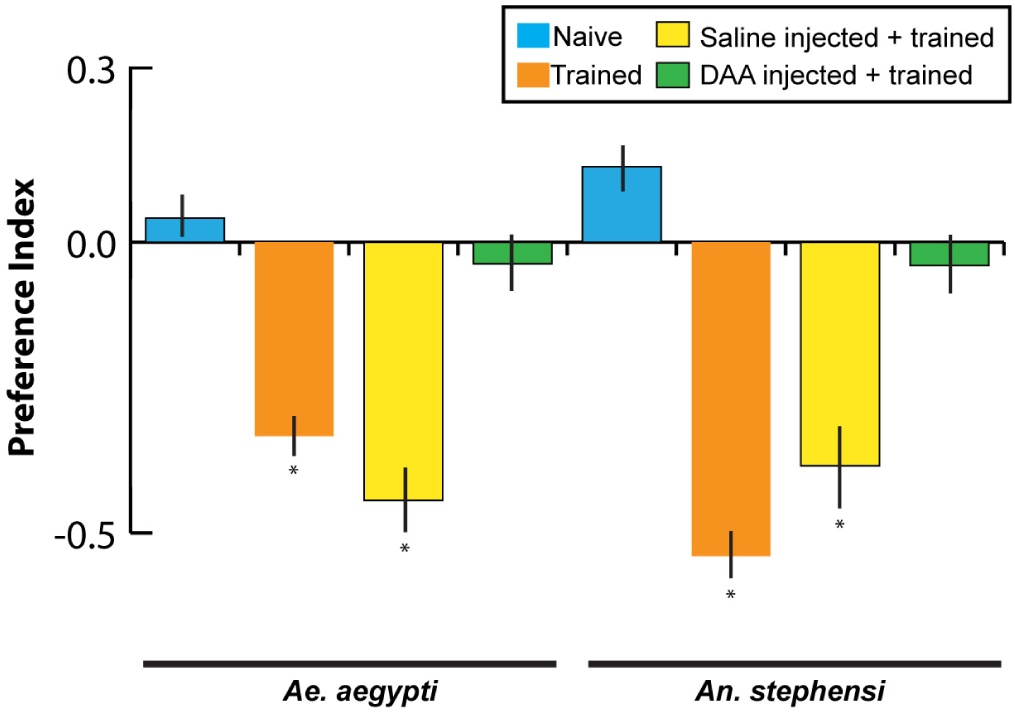
Necessity of dopamine signaling for aversive olfactory learning across species. Preference index of female *Ae. aegypti* and *An. stephensi* mosquitoes for 1-octen-3-ol in naïve (blue), trained (orange), saline-injected + trained (yellow), and D1 dopamine receptor antagonist SCH 23390-injected (DAA) (green) treatment groups. Asterisks represent statistical significance from zero (Binomial exact test: p<0.05). Bars are the mean ± SEM.

### Relationship between odor-evoked responses in the *Ae. aegypti* AL dopaminergic innervation

Learning abilities of different mosquito species and the importance of dopamine in this behavior raised the question of what is the functional relationship between odor-evoked glomerular response and dopaminergic innervation? To understand this relationship, we first sought to map odor-evoked responses of 1-octen-3-ol, hexanoic acid, and linalool in the *Ae. aegypti* AL glomeruli and compare relative concentrations of TH expression between the glomeruli that encode these odors. We used our GCaMP6s *Ae. aegypti* line to record glomerular activity while stimulating mosquitoes with our odors of interest and a mineral oil control (Fig. 5a-g; Extended Data Fig. 2c,d; Extended Data Fig. 3a,d). Serial imaging of depths across the AL – composed of more than 53 glomeruli^17^ – enabled us to analyze 18 glomeruli that were reliably identifiable across preparations based on their position. Some of these glomeruli are responsive to odorants like nonanal, lilac aldehyde and other compounds^19,20^, some of which are not learnable by *Ae. aegypti*. Stimulation with 1-octen-3-ol elicited the largest response in the MD3 glomerulus which receives innervation from the maxillary palps where the 1-octen-3-ol receptor AeaegOR8 is expressed (Kruskal-Wallis test with multiple comparisons: p=0.0019; Fig. 5b, e; ^21^). By contrast, other glomeruli were not significantly different from one another in their responses to 1-octen-3-ol (p>0.05), although their responses (e.g., AM4, PC1) were still significantly greater than mineral oil (no odor) control (p<0.05). Hexanoic acid only elicited significant responses in the MD1 glomerulus (p<0.01) compared to other glomeruli, although other glomeruli were also mildly activated (e.g., V3, PD1, PC1, and PL6; Fig. 5c,f). One glomerulus, AL3 responded significantly to hexanoic acid but also responded robustly to 1-octen-3-ol, linalool, and every other odor we tested and may be very broadly tuned (Extended Data Fig. 5b,c). Similar to hexanoic acid, linalool elicited broad responses across glomeruli, although only the responses by the AM4 glomerulus was significantly higher (p=0.041; Fig. 5d,g).

**Fig. 5.**
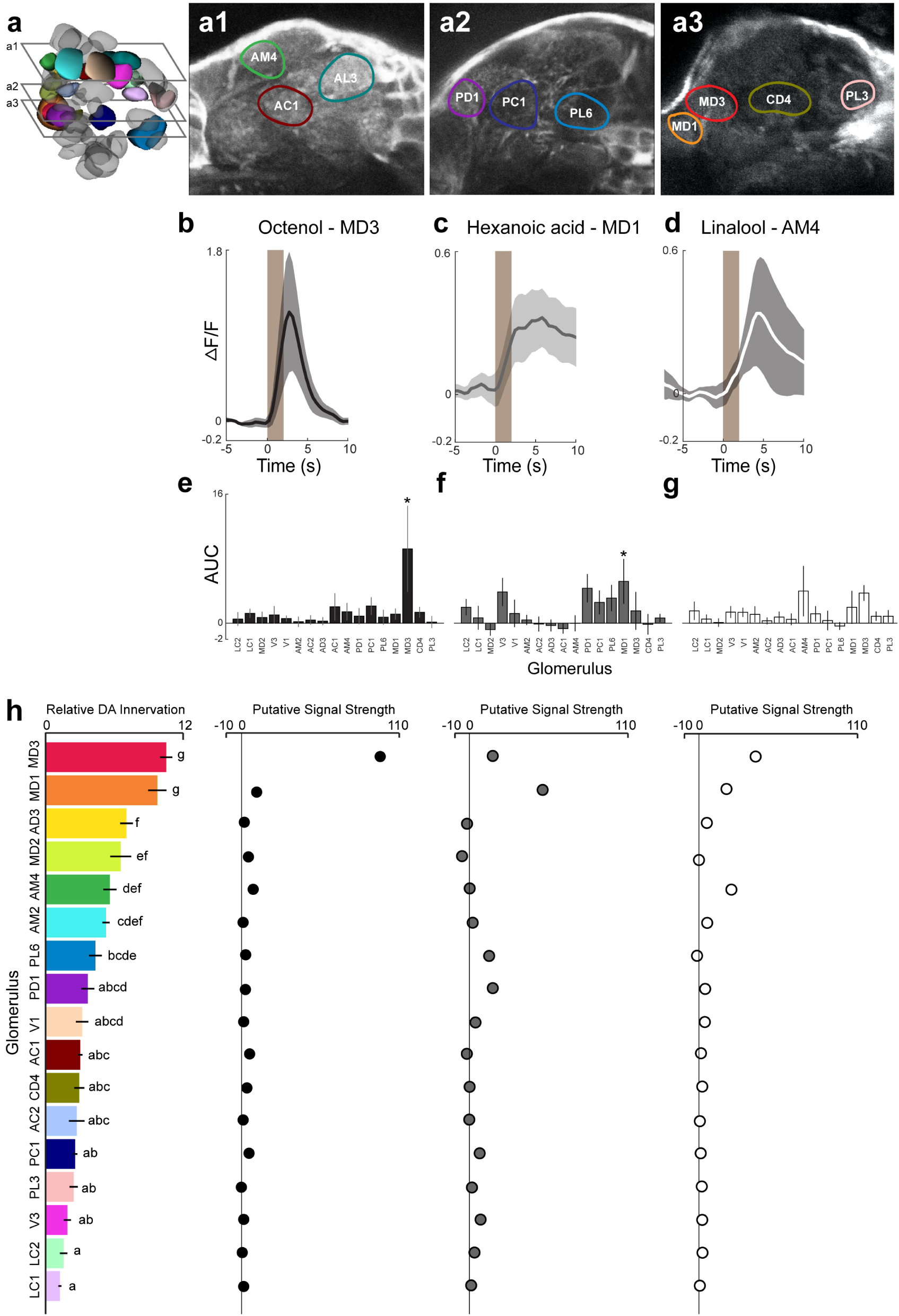
Odor-evoked activity and relative dopaminergic innervation of olfactory glomeruli. **a**, 3-dimensional schematic of the *Ae. aegypti* female antennal lobe. Glomeruli analyzed in this study are highlighted and colors correspond to their labels in h. Representative frames from calcium imaging videos with outlined regions of interest are shown at depths (distances along anterior-posterior axis measured from anterior surface of the antennal lobe) 20-30µm (**a1**), 50-60 µm (**a2**), and 75-80µm (**a3**). Normalized averages of calcium responses (ΔF/F) to (**b**) 1-octen-3-ol in glomerulus MD3, (**c**) hexanoic acid in glomerulus MD1, and (**d**) linalool in glomerulus AM4. e-g, Average area under the curve (AUC) of each calcium response for ten seconds beginning at the time of stimulus onset when mosquitoes were presented with **e** 1-octen-3-ol, **f**, hexanoic acid, and **g**, linalool. Asterisks represent level of significance in a linear regression model. *p<0.01. **h**, Left: Relative dopamine (DA) innervation calculated as normalized average signal to noise from TH-like immunoreactivity (average pixel intensity of glomerulus divided by average pixel intensity of background). Letters represent significance groups (1-factor ANOVA and Tukey post hoc tests, p<0.05). Right: Putative strength of signal output (due to neuromodulation) from each glomerulus as a function of relative DA innervation multiplied by calcium response (AUC) to 1-octen-3-ol (black), hexanoic acid (grey), and linalool (white).

We next examined the relationship between dopaminergic innervation and glomerular responses to odorants. Dopaminergic innervation was mapped to each glomerulus by staining for TH and calculating the signal to noise (average pixel intensity within the glomerulus divided by average pixel intensity of the background) correlated with odorant-evoked responses across the glomerular ensemble (Fig. 5h; Extended Data Fig. 1c). When we compared the relationship between 1-octen-3-ol responses and TH levels across glomeruli, we found a significant relationship between 1-octen-3-ol responses and dopamine levels (linear model: r=0.52; F1,15=5.68; p=0.03), with MD1 responses also reflected in the strong dopaminergic innervation (p<0.05). Similarly, the strong calcium-evoked responses by MD3 to hexanoic acid were also reflected in the significant dopaminergic innervation to this glomerulus (p<0.01), with 1.7 to 7.3-fold higher innervation than other glomeruli. By contrast, for linalool, there was no relationship between the glomerular responses and dopamine levels (p=0.25). To examine how this innervation might filter the input to downstream circuits in the MB, we convolved the odor-evoked response and the TH levels to produce a putative signal strength for each glomerulus (Fig. 5h). Results showed that for both 1-octen-3-ol and hexanoic acid, MD1 and MD3 glomeruli were amplified by 110- and 63-fold higher levels, respectively, over other glomeruli, whereas for glomeruli activated by linalool (e.g., AM4, MD1 and LC2), the signal strength was significantly lower (mean=11.8 ± 2.5; Kruskal-Wallis test with multiple comparisons: p<0.05; Fig. 5h), together indicating that the glomeruli encoding odors learned by *Ae. aegypti* had high levels of dopaminergic innervation relative to other glomeruli.

## Discussion

Olfactory learning has long been implicated in shaping insect behaviors when preferred resources are scarce, however, the degree of learning by different mosquito species and its relationship to their host preferences has remained an open question. Here, we show that learning preferred-host odors may be a common phenomenon across mosquitoes, and that dopamine plays an important role in mediating this process. Furthermore, using *Ae. aegypti*, our results suggest a relationship between dopaminergic innervation to specific antennal lobe glomeruli and a mosquito’s ability to learn the odor encoded therein. Dopaminergic modulation in olfactory learning has primarily been studied in the mushroom bodies^22–24^, which receive afferents from the ALs and integrate signals from different senses as well as reward and punishment. However, neuromodulation at the level of the AL can induce neuroplasticity, mediate gain control of olfactory signals, and change the strength of output to higher-order centers like the mushroom bodies^25^. In *Ae. aegypti*, dopamine has differential modulatory effects on glomeruli encoding different odors. For example, representation of 1-octen-3-ol in the AL was significantly changed upon application of dopamine, increasing its discriminability with respect to other stimuli^4^. Here we show that glomeruli tuned to odors learned by mosquitoes, such as 1-octen-3-ol, are highly innervated with dopaminergic fibers compared to other glomeruli. The specific identities of modulatory neurons innervating glomeruli are yet to be determined, although improved genetic tools may allow us to functionally image and interrogate their role in AL modulation and downstream effects on MB Kenyon cells.

This study also highlights the importance of a comparative approach to understanding mosquito learning and food preferences. Our results suggest those species that prefer to feed on human blood aversively learned odors associated with mammalian skin volatiles but not a flower odor and the opposite was true of mosquitoes that feed solely on plant carbohydrates. Although the number of tested odorants in our behavioral and electrophysiological assays was relatively small, the odorants evoked similar levels of afferent drive to the antennal lobe and reflected the odorants from different nutrient resources the mosquitoes might utilize. *Cx. quinquefasciatus* did not learn in our classical conditioning paradigm with an aversive stimulus designed to replicate host defensive behaviors. One explanation may be because of the lack of dopaminergic innervation in the AL, or alternately, that because this species is nocturnal (while the others are diurnal and crepuscular), host animals are more likely to be sleeping at peak activity hours and less likely to display defensive behaviors^5,26^. Thus, there may be less selective pressure on *Cx. quinquefasciatus* to evolve aversive learning circuitry.

Evolution of neuromodulatory inputs to sensory and motor circuits played a role in mediating species-specific differences in behavior across animal phyla^27^. Our study suggests that evolution of species-specific host preferences in mosquitoes may have co-evolved with differences in dopaminergic innervation to the antennal lobe. For all four species, patterns of dopaminergic innervation varied, although the basic morphology and circuit organization is conserved. Indeed, mosquitoes share ground pattern organization of olfactory circuitry with all insects and glomerular structure is conserved across invertebrate olfactory lobes and vertebrate olfactory bulbs^28^. An evolved increase in dopaminergic input to certain glomeruli may prime those circuits to relay odor-specific information to higher-order centers of learning and memory. When host availability changes on a short evolutionary timescale, mosquitoes may adapt to learning new olfactory signals by modifying dopaminergic inputs to prime the most salient odor circuits. Dopaminergic systems are involved in arousal, developmental processes, and results from this study suggest flexibility in host preferences^29,30^. Moreover, olfactory behaviors, including learning, are critical for mosquito preferences and biting of human hosts, and thereby the spread of diseases that afflict over a billion people annually. Therefore, unraveling the neural basis of learning in mosquitoes provides motivation for the development of new strategies for their control.

## Methods

### Animals

*Aedes aegypti* (wild type: MRA-734, ATCC®, Manassas, VA, USA; and our PUb-GCaMP6s line), *An. stephensi* (MRA-128, Strain STE2, CDC, Atlanta, GA, USA) and *Cx. quinquefasciatus* (NR-43025, Strain JHB, CDC, Atlanta, GA, USA) were maintained in breeding colonies at the University of Washington. Toxorhychites amboinensis eggs were a generous donation from Jason Pitts (Baylor University, Waco, Texas) and were also used to establish a new breeding colony. Mosquitoes were reared at 25±1°C, 60±10% relative humidity (RH) in climatic chambers that maintained a 12 hour light cycle (12:12). *Ae. aegypti, An. stephensi* and *Cx. quinquefasciatus* mosquitoes were fed heparinized bovine blood (Lampire Biological Laboratories, Pipersville, PA, USA) through a parafilm membrane attached to a double-walled glass feeder (D.E. Lillie Glassblowers, Atlanta, GA, USA; 2.5 cm internal diameter) circulating water at 37C. All species were provided with 10% sucrose. Eggs were hatched in trays of deionized water and larvae were provided with powdered fish food (Hikari Tropic 382 First Bites - Petco, San Diego, CA, USA). *Tx. amboinensis* eggs were hatched in cups of deionized water and larvae were placed in separate containers of deionized water (to prevent cannibalism) and fed *Ae. aegypti* larvae. Pupae of all species were collected and placed in mosquito breeding jars and, once eclosed, they were maintained on 10% sucrose (Bio-Quip, Rancho Dominguez, CA, USA). Mosquitoes used in all experiments were given a chance to mate, but were non-blood fed and isolated at six days post-eclosure.

### Antibodies

Whole brains were double-labeled with mouse monoclonal antisera against tyrosine hydroxylase (ImmunoStar, Hudson, WI, USA - Cat. no. 22941) used at a concentration of 1:50 and rabbit polyclonal antisera against phosphorylated CaMKII (Santa Cruz Biotechnology, Dallas, TX, USA - Cat. no. sc-12886R) were used at a concentration of 1:100 for immunohistochemistry. Sectioned brains were double-labeled with mouse monoclonal antisera against glutamine synthetase (BD Bioscience, Cat. no. 610517), used at a concentration of 1:200 and rabbit polyclonal antisera against GFP (abcam, Ca. no. ab6556) were used at a concentration of 1:500.

### Immunohistochemistry

Mosquitoes were placed in a refrigerator at 4 C until immobile and heads were dissected into 4 C fixative containing 4% paraformaldehyde in phosphate-buffered saline, pH 7.4 (PBS, Sigma-Aldrich, St. Louis, MO, USA -Cat. No. P4417). After one hour in fixative, brains used for tyrosine hydroxylase and pCaMKII immunohistochemistry were placed into PBS containing 4% Triton X-100 (PBS-TX; Sigma-Aldrich, St. Louis, MO, USA - Cat. No. X100) and incubated in that solution overnight. Next, brains were placed in glass vials and washed two times for 10 minutes each in 0.5% PBS-TX. 50 µL normal serum and 1,000 µL 0.5% PBS-TX was added to each vial and incubated for one hour. Then primary antibody was added to each vial and the vials were placed on a shaker for 48 hours at 4C. Next, brains were washed six times for 20 minutes each in PBS-TX. 1,000-µL aliquots of PBS-TX with 2.5 µL of secondary Alexa Fluor 488 or Alexa Fluor 546-conjugated IgGs (ThermoFisher Scientific, Waltham, MA, USA) were centrifuged at 13,000 rpm for 15 minutes. The top 900-µL of this solution was added to each glass vial and brains or sections were incubated in this solution for 48 hours on a shaker at 4C. Brains were next washed in PBS six times for 20 minutes each and embedded on glass slides in a medium containing 25% polyvinyl alcohol, 25% glycerol, and 50% PBS. For GFP and glutamate synthetase staining, brains were first embedded in agarose and sectioned at 60µm on a vibrating microtome before following the above steps, with the modification that primary and secondary antibody incubations were 24 hours at room temperature. Immunofluorescence was imaged on a Leica SP5 laser scanning confocal microscope (Leica Microsystems, Wetzlar, Germany). Image stacks were processed as maximum intensity projections using ImageJ (National Institutes of Health) and selected images were assembled using Adobe Photoshop CS4 (Adobe Systems, San Jose, CA, USA).

### Image Analysis

Tyrosine hydroxylase immunofluorescence from confocal image stacks was quantified using ImageJ software. The Segmentation Editor plugin was first used to reconstruct glomeruli in 3D. For each of these volumes, average pixel intensities were calculated using the 3D Manager plugin. To correct for shielding effects, average pixel intensities were calculated in the same manner for volumes of background signal at the depth of each glomerulus. “Relative TH concentration” was calculated as average pixel intensity within a glomerulus divided by average pixel intensity within a spherical volume, 8µm in diameter at the same depth as the glomerulus but located in an area of background.

### Electro-antennogram (EAGs)

Mosquito heads were excised and the tip of each antenna (i.e. about one segment) was cut off with fine scissors under a binocular microscope (Carl Zeiss, Oberkochen, Germany). The head was mounted on a borosilicate pulled capillary (Sutter Instrument Company, Novato, CA, USA) with an inserted silver wire 0.01” (A-M Systems, Carlsbord, WA, USA) and filled with a 1:3 mix of saline solution (Beyenbach and Masia, 2002) and electrode gel (Parker Laboratories, Fairfield, NJ, USA); this electrode served as the reference. The preparation was then moved to the EAG setup and using a micromanipulator (Narishige, Japan), the tips of the antennae were inserted under the microscope (Optiphot-2, Nikon, Tokyo, Japan) into a recording electrode, identical to the reference electrode. The mounted head was oriented at 90° from the main airline which was carrying clean air (Praxair, Danbury, CT, USA) to the preparation for the whole duration of the experiment. Odor stimuli were injected into the airstream via a computer-triggered solenoid valve. Each odor was pulsed twice for three seconds with a ten second inter-stimulus interval and EAG responses were recorded using the software WinEDR (V3.8.6, Strathclyde Electrophysiology Software; Strathclyde University, Gasglow, UK). Responses were calculated in R (www.r-project.org) as the amplitude from peak to trough.

### 2-photon Calcium Imaging

Odor-evoked responses were imaged in the AL of female *Ae. aegypti* expressing genetically-encoded polyubiquitin-GCaMP6s during tethered flight^31^. Calcium responses were visualized on a Prairie Ultima IV multiphoton microscope (Prairie Technologies) and Ti-Sapphire laser (Chameleon Ultra, Coherent). Laser power was adjusted to 20mW at the rear aperture of the objective lens (Nikon NIR Apo, 40X water immersion lens, 0.8 NA), GCaMP fluorescence was bandpass filtered with a HQ 525/50 m-2p emission filter (Chroma Technologies) and photons collected using a multialkali photomultiplier tube. Individual mosquitoes were cold-anaesthetized on ice and transferred to a Peltier-cooled holder while the head was glued to a 3D-printed stage. This custom stage is shaped like an inverted pyramid with a window at the vertex to permit superfusion of saline to the head capsule and room for the mosquito to move its abdomen and beat its wings. After gluing the head to the stage, a window was cut in the cuticle to expose the brain and a cold, oxygenated saline drip was placed in the well. A continuous stream of filtered air was directed towards the mosquito’s antennae and odors were injected through an odor cartridge into this stream via solenoid valve for 2 seconds per stimulation. Odor cartridges consisted of glass syringes containing filter paper and 2µL of either mineral oil or odor chemical (1-octen-3-ol, hexanoic acid, or linalool) diluted in mineral oil at 1:100. For each odor stimulation, images of the antennal lobe were collected at 2 Hz over 30 seconds, beginning 10 seconds before odor stimulus onset. These images – 500 µm by 600 µm window, sampled at 2 Hz – were time-stamped and synchronized with the time course of the odor-stimulus. Calcium responses were collected while imaging at focus planes 20-30 µm, 50-60 µm, or 70-80 µm from the top of the antennal lobe in the anterior-posterior neuraxis.

Images were examined in Fiji and if two-dimensional movement occurred, the images were exported to Matlab (v2018; Mathworks, Natick, Massachusetts) for Gaussian filtering (2×2 pixel; σ = 1.5-3) and alignment using a single frame as the reference at a given imaging depth and subsequently registered to every frame to within ¼ pixel. Glomerular ROIs were determined based on the clear boundaries around glomeruli, especially during odor stimulation. We also note that the GCaMP6s is expressed in multiple cell types in the *Ae. aegypti* brain and AL, including glia, projection neurons, local interneurons, and olfactory sensory neurons. Glia appeared in astroglial-like projections around the exterior of the glomeruli, whereas during odor stimulation dendrites and axons from projection neurons and local interneurons within the interior of the glomeruli often become apparent. Odor-evoked calcium responses were calculated as the change in fluorescence for each glomerulus in each frame t of a time series using the formula: ΔF/F_0_ = (F_t_ −F_0_)/F_0_, where F_0_ is the average intensity during the baseline period prior to the presentation of the odor stimulus. For each odor, the area under the curve (AUC) – reflecting the total response of the glomerulus and calculated by summing the ΔF/F for each frame from the stimulus onset to the signal return to baseline – was calculated for each glomerulus. We used the glomerular AUC for most statistical tests as this may reflect the total input to the MB. Finally, optical sections (1 µm) through the AL were collected with the same multiphoton microscope and olfactory glomeruli were reconstructed in 3D using ImageJ and the Segmentation Editor plugin.

### Behavioral Assays

#### Mosquito training protocol and control groups

A total of 1,944 individual female mosquitoes were used in the behavioral experiments. Aversive conditioning was performed as in Vinauger et al., 2018 (Extended Data Fig. 1b). Briefly, female mosquitoes were individually separated into plastic containers with mesh openings and allowed to acclimate for 120 seconds in a training chamber with a tube delivering clean (medical grade) air (30 cm.s-1, 23° C, 50% RH). Then, an odor was delivered via solenoid valve into a separate air stream for 60 seconds. During the last 30 seconds of odor presentation, the odor was paired with a mechanical shock delivered by a vortexer (1.65 g at 44 Hz). Odor and mechanical shock pairings were presented ten times with a 2-minute inter-trial interval (ITI). During the ITI, odor was vacuumed out of the training chamber while the clean airline continued to flow. After the training session was completed, mosquitoes were placed in a climatic chamber (25° C; 60% RH; 12-12 h L:D) for 24 hours before testing in a Y-maze olfactometer. Both training and testing occurred at 10 hours into the light phase of the 12:12 light cycle for *Ae. aegypti* and *An. stephensi*, and at 8 hours for *Tx. amboinensis. Cx. quinquefasciatus* were trained and tested at the onset of the dark phase in the 12:12 light cycle. These time periods reflect the peak activity time of the mosquitoes. Two control groups were compared to the trained group to test the effects of aversive conditioning. A naïve group was tested for each species and in cases when preference of trained mosquitoes to an odor differed significantly from naïve, an unpaired group was tested as well. Unpaired mosquitoes were placed in the training chamber and handled in the same manner as trained mosquitoes except that odor and mechanical shock were delivered semi-randomly, without overlap. Each group was placed in the climatic chamber for 24 hours before testing. In previous studies, conditioned-stimulus-only and unconditioned-stimulus-only groups did not behave significantly different from naïve groups in this paradigm4.

Training and testing trials were conducted in blocks of 12 or 24 mosquitoes. Certain groups were tested throughout the study period to validate results and therefore have higher sample sizes. These include ‘unpaired’ *Ae. aegypti* tested against 1-octen-3-ol, naive *Ae. aegypti* tested against linalool, and trained *Cx. quinquefasciatus* tested against hexanoic acid.

#### Behavioral testing in the olfactometer

Odor preference was tested using a custom-built, Plexiglas® Y-maze olfactometer as described by Vinauger et al., 2018. Mosquitoes were placed in a starting chamber attached to a cylindrical entry tunnel (30 cm long, 10 cm diameter), leading to a central chamber connected to two symmetrical “choice” tunnels (both 39 cm long, and 10 cm diameter). Fans drew air into each “choice” arm through a charcoal filter and a honeycomb filter to create uniform laminar flow at at 20 cm.sec-1. Odor and control stimuli were delivered by pumping air into the “choice” tunnels via Teflon® tubing connected to one of two 20 mL scintillation vials containing 10 mL of either mineral oil (control) or tested odor (1-octen-3-ol 1:100, hexanoic acid 1:100, linalool 1:1000 diluted in mineral oil). Control and tested odor presentation were randomized daily between the left and right “choice” arms and the maze was cleaned with 70% and 100% ethanol after each experiment. Behavioral testing was performed in a climatic chamber at 27°C and 70% RH.

Each test consisted of placing a single female mosquito in the starting chamber of the olfactometer, then the experimenter exited the climatic chamber and simultaneously monitored and filmed the maze via webcam. The first choice of the mosquito was recorded when she crossed the entry of a “choice” arm. Mosquitoes that did not make a choice within 5 minutes were marked as active and non-responsive if they flew as far as the central chamber and they were marked as inactive and non-responsive if they did not leave the starting chamber. These two non-responsive groups were discarded from the preference analysis. Activity and percent choice for each species and group is reported in Table S1.

### Statistical analyses

Analyses were performed in R. For the binary data collected in the olfactometer, comparisons were performed by means of the Binomial Exact test (α=0.05). For each treatment, the choice of the mosquitoes in the olfactometer was either compared to a random distribution of 50% on each arm of the maze or to the distribution of the corresponding control when appropriate. For binary data, the standard errors (SE) were calculated: SEM =((p(1-p))/n)^(1/2) For each experimental group, a preference index (PI) was computed in the following way: PI = [(number of females in the test arm – number of females in the control arm) / (number of females in the control arm + number of females in the test arm)]. A PI of +1 indicates that all the motivated mosquitoes chose the test arm, a PI of 0 indicates that 50% of insects chose the test arm and 50% the control arm, and a PI of -1 indicates that all insects chose the control arm of the olfactometer. For the calcium imaging, EAG and IHC data, whenever the data did not meet the normality assumption of the ANOVA, a Kruskal-Wallis test (with post hoc Tukey test for multiple comparisons) was performed. A linear regression model was used to examine the relationship between glomerular responses and TH-innervation.

## ACKNOWLEDGEMENTS

We thank Benjamin Arnold, Dane Alzate, Brian Vickers, and Erica Peterson for performing olfactometer experiments. We also thank William Moody, Tom Daniel, Claire Rusch, Diego Alonso San Alberto and Tom Libby for coding and technical assistance. *Aedes aegypti, Anopheles stephensi*, and *Culex quinquefasciatus* photographs are credited to James Gathany and the Center for Disease Control. We acknowledge the support of the Air Force Office of Scientific Research under grant FA9550-16-1-0167, National Science Foundation under grant 1354159; National Institutes of Health under grant 1R21AI137947, an Endowed Professorship for Excellence in Biology, the University of Washington Innovation Award.

## Author contributions

G.H.W., C.L., C.V., and J.A.R conceived and designed the experiments; G.H.W. and C.L. performed the experiments; G.H.W. and J.A.R. analyzed the data; all authors contributed to the writing and editing of the manuscript and all authors read and approved the final manuscript.

## Extended Data

**Extended Data Figure 1.**
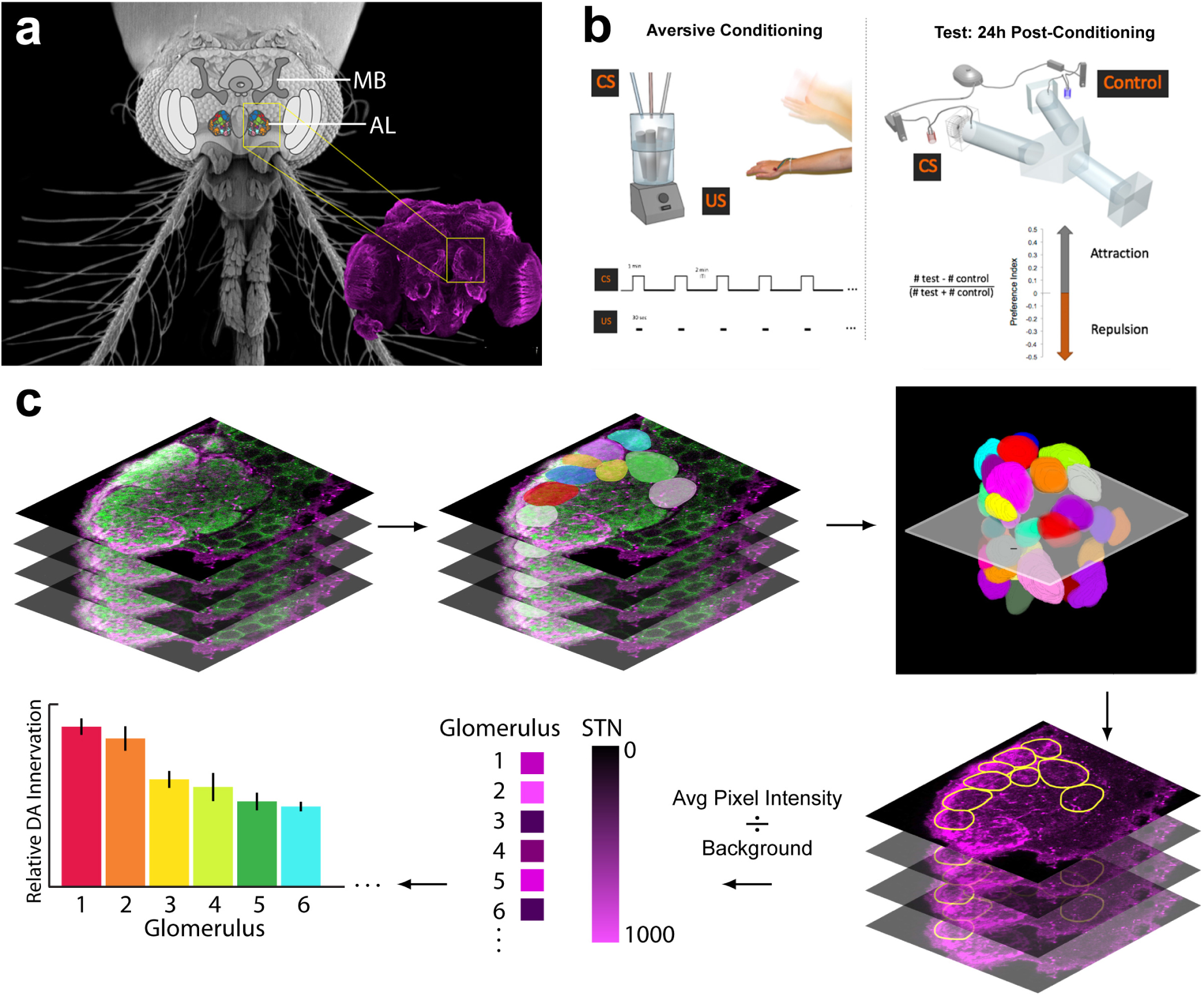
Experimental design. **a**, Schematic of a mosquito brain superimposed on scanning electron micrograph (provided by Joseph Dickens, USDA-ARS) of an *Ae. aegypti* female head to show relative position. Inset: *Ae. aegypti* brain stained with antisera against alpha-tubulin. Yellow boxes indicate position of right antennal lobe (AL). MB= mushroom bodies. **b**, Aversive classical conditioning paradigm. Left: Mosquito training chamber and timing of unconditioned (US) and conditioned (CS) stimuli presentation. Right: Y-maze olfactometer for testing mosquito preference. **c**, Workflow for calculating relative dopaminergic innervation. Z-stacks of antennal lobes from 5 individual female *Ae. aegypti* stained with antisera against tyrosine hydroxylase (magenta) and CaMKII (green) were collected on a confocal microscope. Glomeruli were traced through each stack and then reconstructed in 3D. Each 3D glomerulus volume was used as a region of interest overlaid on the tyrosine hydroxylase channel. Using the “3D Manager” plugin in ImageJ, average pixel intensity was calculated for each region of interest. This value was divided by the background at the depth of each glomerulus to calculate signal to noise (STN) which served as a proxy for relative dopamine innervation.

**Extended Data Figure 2.**
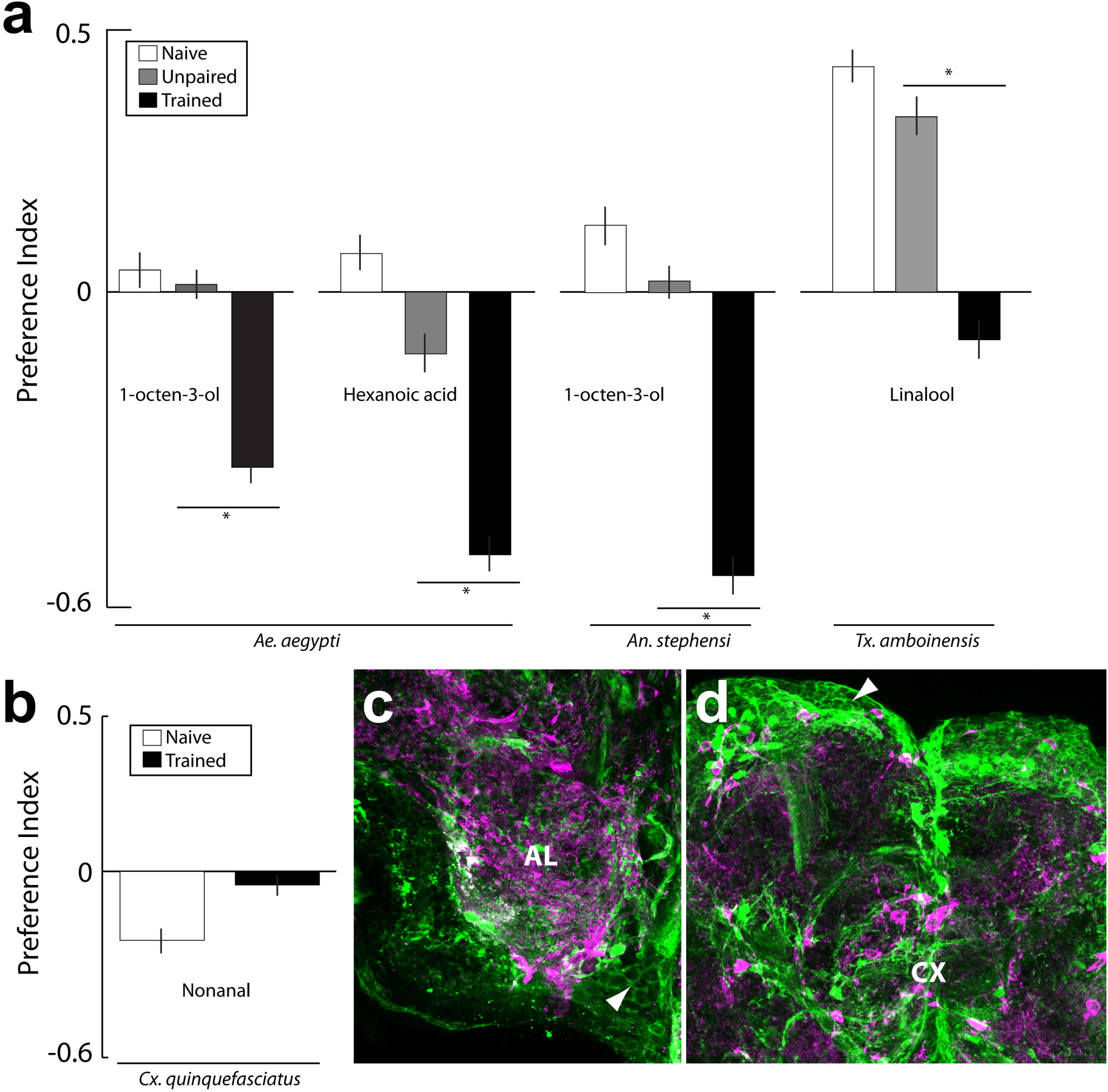
Unpaired and imaging controls. **a**, To control for the effects of aversive conditioning, trained and naïve mosquitoes were compared to an “Unpaired” control group in cases where the former groups differed significantly in preference. Trained *Ae. aegypti, An. stephensi*, and *Tx. amboinensis* differed significantly from unpaired, while unpaired were not significantly different from naïve (* p<0.05, Binomial exact test). **b**, Preference of female *Cx. quinquefasciatus* to nonanal in trained and naïve groups were not significantly different. Bars are the mean ± SEM. **c-d**, Confocal laser scans of PUb-GCaMP6s *Ae. aegypti* female brains, sectioned and stained with antisera against GFP (green) and glutamate synthetase (magenta). AL = antennal lobe, CX = central complex. Arrowheads indicate neuron cell bodies expressing GFP.

**Extended Data Figure 3.**
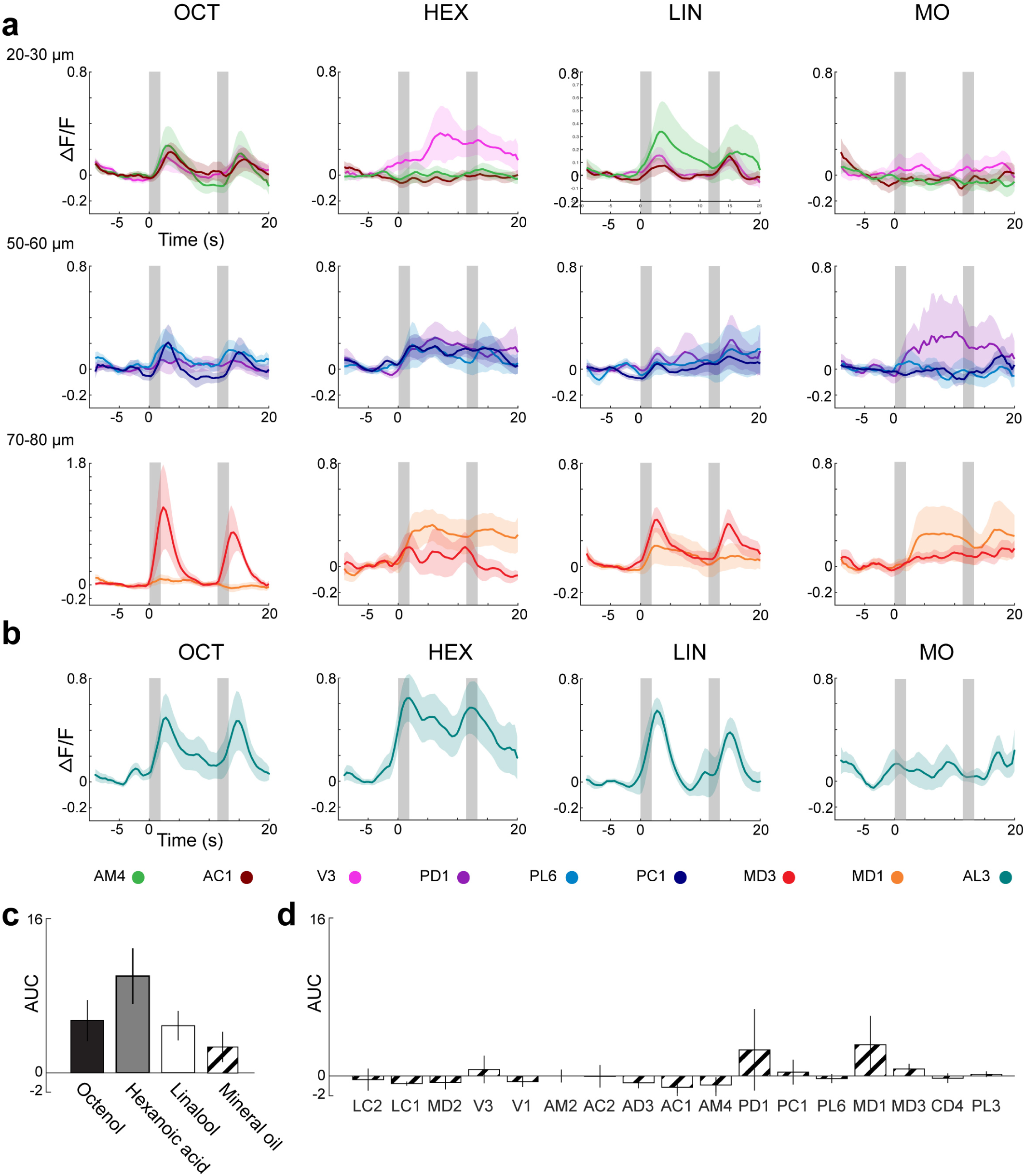
Odor evoked calcium responses. **a**, Normalized averages of glomerular responses to 1-octen-3-ol (OCT), hexanoic acid (HEX), linalool (LIN), and mineral oil (MO) at three different focal planes (distances along anterior-posterior axis measured from anterior surface of the antennal lobe). Colors correspond to glomeruli identities labeled in Fig. 5h. **b**, The AL3 glomerulus was activated by all tested odorants and appears to respond generally to a broad range of odor stimuli, including the mineral oil control. **c**, Calcium response for AL3 was quantified as the area under the curve (AUC) for ten seconds after the time of stimulus onset. **d**, AUC quantified for the remaining glomeruli in response to mineral oil.

**Extended Data Table 1:**
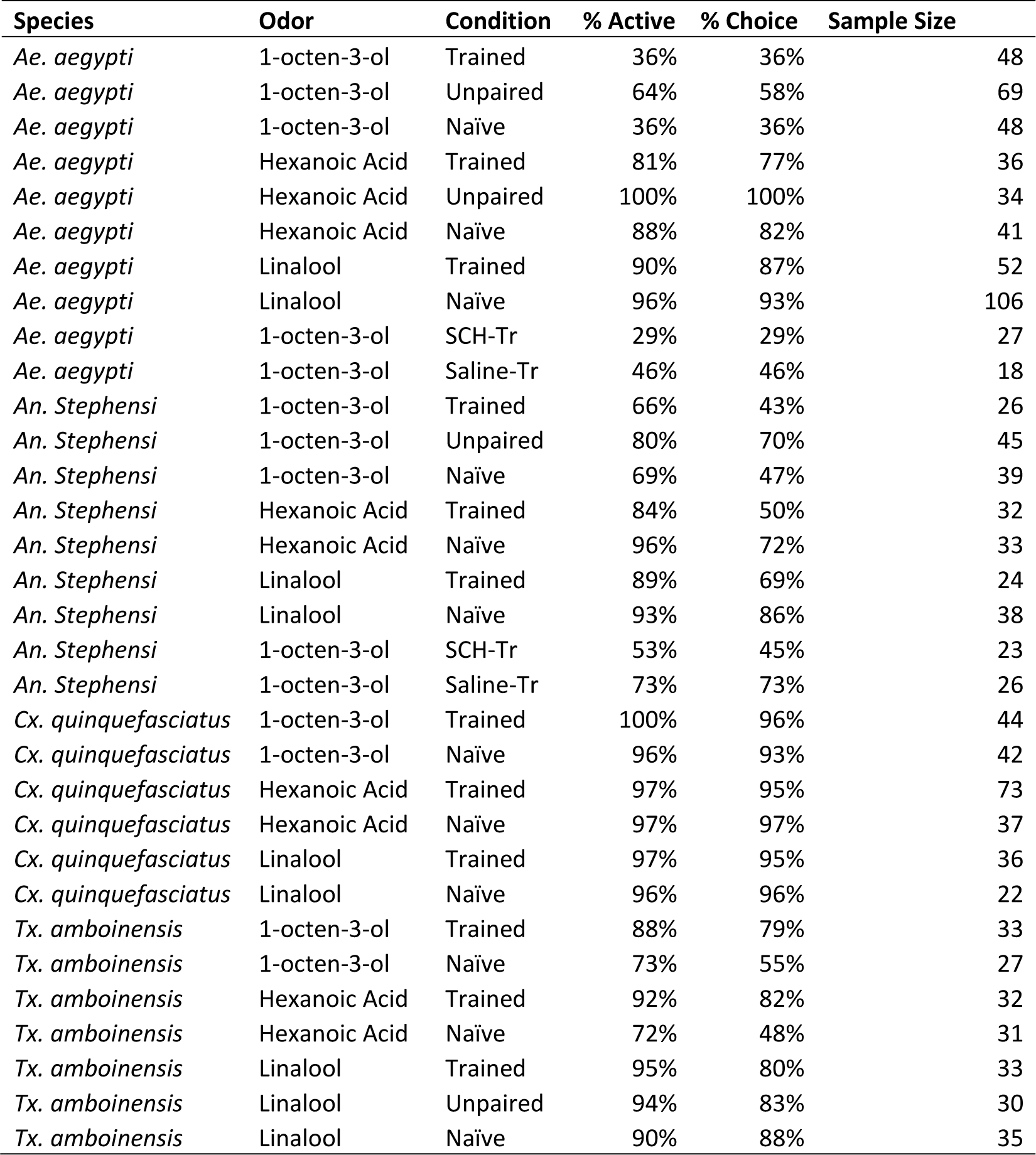
Activity levels of mosquitoes in behavioral assays. %Active represents all mosquitoes in the group who either flew to the central chamber or down one of the “choice arms” in the olfactometer. %Choice represents only those mosquitoes that flew down a “choice” arm as a percentage of the total. SCH-Tr = mosquitoes injected with the dopamine antagonist, Saline-Tr = saline-injected controls. Sample Size includes only those mosquitoes that made a choice.

| Species | Odor | Condition | % Active | % Choice | Sample Size |
| --- | --- | --- | --- | --- | --- |
| *Ae. aegypti* | 1-octen-3-ol | Trained | 36% | 36% | 48 |
| *Ae. aegypti* | 1-octen-3-ol | Unpaired | 64% | 58% | 69 |
| *Ae. aegypti* | 1-octen-3-ol | Naïve | 36% | 36% | 48 |
| *Ae. aegypti* | Hexanoic Acid | Trained | 81% | 77% | 36 |
| *Ae. aegypti* | Hexanoic Acid | Unpaired | 100% | 100% | 34 |
| *Ae. aegypti* | Hexanoic Acid | Naïve | 88% | 82% | 41 |
| *Ae. aegypti* | Linalool | Trained | 90% | 87% | 52 |
| *Ae. aegypti* | Linalool | Naïve | 96% | 93% | 106 |
| *Ae. aegypti* | 1-octen-3-ol | SCH-Tr | 29% | 29% | 27 |
| *Ae. aegypti* | 1-octen-3-ol | Saline-Tr | 46% | 46% | 18 |
| *An. Stephensi* | 1-octen-3-ol | Trained | 66% | 43% | 26 |
| *An. Stephensi* | 1-octen-3-ol | Unpaired | 80% | 70% | 45 |
| *An. Stephensi* | 1-octen-3-ol | Naïve | 69% | 47% | 39 |
| *An. Stephensi* | Hexanoic Acid | Trained | 84% | 50% | 32 |
| *An. Stephensi* | Hexanoic Acid | Naïve | 96% | 72% | 33 |
| *An. Stephensi* | Linalool | Trained | 89% | 69% | 24 |
| *An. Stephensi* | Linalool | Naïve | 93% | 86% | 38 |
| *An. Stephensi* | 1-octen-3-ol | SCH-Tr | 53% | 45% | 23 |
| *An. Stephensi* | 1-octen-3-ol | Saline-Tr | 73% | 73% | 26 |
| *Cx. quinquefasciatus* | 1-octen-3-ol | Trained | 100% | 96% | 44 |
| *Cx. quinquefasciatus* | 1-octen-3-ol | Naïve | 96% | 93% | 42 |
| *Cx. quinquefasciatus* | Hexanoic Acid | Trained | 97% | 95% | 73 |
| *Cx. quinquefasciatus* | Hexanoic Acid | Naïve | 97% | 97% | 37 |
| *Cx. quinquefasciatus* | Linalool | Trained | 97% | 95% | 36 |
| *Cx. quinquefasciatus* | Linalool | Naïve | 96% | 96% | 22 |
| *Tx. amboinensis* | 1-octen-3-ol | Trained | 88% | 79% | 33 |
| *Tx. amboinensis* | 1-octen-3-ol | Naïve | 73% | 55% | 27 |
| *Tx. amboinensis* | Hexanoic Acid | Trained | 92% | 82% | 32 |
| *Tx. amboinensis* | Hexanoic Acid | Naïve | 72% | 48% | 31 |
| *Tx. amboinensis* | Linalool | Trained | 95% | 80% | 33 |
| *Tx. amboinensis* | Linalool | Unpaired | 94% | 83% | 30 |
| *Tx. amboinensis* | Linalool | Naïve | 90% | 88% | 35 |

